# SARS-CoV-2 causes brain inflammation via impaired neuro-immune interactions

**DOI:** 10.1101/2022.07.13.499991

**Authors:** Naomi Oka, Kazuya Shimada, Azusa Ishii, Nobuyuki Kobayashi, Kazuhiro Kondo

## Abstract

The brain inflammation that frequently occurs in SARS-CoV-2 is the cause of neurological complications^1,2^ and long COVID ^3,4,5^. However, many aspects of its pathogenesis mechanism remain unknown ^6, 7^ and no method of treatment has been established ^8^. By administering a non-proliferating adenovirus vector expressing SARS-CoV-2 S1 protein into the nasal cavity of mice, we developed a mouse model (S1 mouse) reproducing brain inflammation, fatigue, depressive symptoms, and lung inflammation. Having intracellular calcium elevating activity, S1 protein increased olfactory bulb apoptosis, and reduced the number of acetylcholine producing cells in the medial septal and the diagonal band of Broca as well as the amount of acetylcholine in the brain. This resulted in disrupting the cholinergic anti-inflammatory pathway (CAP) ^9^ and enhancing inflammation in the brain. Previously, nothing was known about anti-inflammatory factors in the CAP but we discovered that, in the inflammation occurring in the S1 mouse brain, the action of the RNA binding protein ZFP36 ^10^ in degrading inflammatory cytokine mRNA was impaired.

The symptoms exhibited by the S1 mouse were improved by administering donepezil, a drug with a cholinergic action used in the treatment of dementia. These findings clarify the mechanism of brain inflammation in COVID-19 and indicate the possibility of applying donepezil in the treatment of neurological complications in COVID-19 and long COVID.

## Main

Severe acute respiratory syndrome coronavirus 2 (SARS-CoV-2) is the causative virus of coronavirus disease 2019 (COVID-19). Mainly affecting the lungs, it causes pneumonia and acute respiratory distress syndrome (ARDS). Also affecting the kidneys, brain, heart, liver and other organs, it can cause multi-organ failure ^11^. Research on the etiology of SARS-CoV-2 has uncovered many neurological complications but many aspects of their pathogenesis mechanisms remain unknown ^6,7^.

Highly prevalent neurological complications are those due to brain inflammation, such as fatigue and depressive symptoms ^1,2^. If they persist, long COVID ^3-5^ may develop, for which there is no established treatment ^8^.

Inflammation and damage in the brain may be caused in 2 different ways, via encephalitis or encephalopathy. In the former, the inflammation and damage are due to proliferation of SARS-CoV-2 in the brain. In the latter, the inflammation and damage occur in the absence of virus proliferation, and are due to the effects of cytokines. While evidence of small amounts of SARS-CoV-2 infection has been seen in the brain in some postmortem case resports^12,13^, many studies have found encephalopathy with no viral proliferation in the brain ^14-17^.

Small animal models are useful in COVID-19 research. However, in some models of SARS-CoV-2 infection, inflammation occurred with viral proliferation in the brain ^18^, or long term observation was not possible due to pneumonia, ^19^ so they have not been suitable disease models for neurological complications.

Therefore, in the present research, our plan was to create a model in which disease was caused by inducing expression of SARS-CoV-2 pathogenic protein in the respiratory tract. Since it has been reported that S1 protein^20,21^ may have an inflammation enhancing function in the brain, we decided to create a model mouse expressing S1 protein.

### Characterization of S1 Expressing Mice

Our objective in creating the model mouse expressing S1 protein (S1 mouse) was to achieve an animal model capable of reproducing the brain inflammation, fatigue, depressive symptoms and pneumonia frequently seen in COVID-19. We prepared a non-proliferative adenovirus vector (S1 Adv) expressing the S1 protein of the original Wuhan strain (Wu strain) and then created the S1 mouse through inoculation of S1 Adv into the nasal cavity (Fig. 1a). One week after inoculation, we observed enhanced expression of inflammatory cytokines (IL-6, TNFα) and a chemokine (CCL-2) with an inflammation promoting function (Fig. 1b, Extended Data Fig.1a, b) suggesting that inflammation had occurred in the brain. Owing to a decrease in swimming time in the weight-loaded forced swim test (WFST) ^22^, we determined that fatigue had increased in the S1 mouse (Fig. 1c). In addition, there was an increase in immobility time for the S1 mouse in the tail suspension test (TST) (Fig. 1d), which we interpreted as presence of depressive symptoms. We also observed enhanced expression of IL-6 in the lungs (Fig. 1e).

**Fig. 1:**
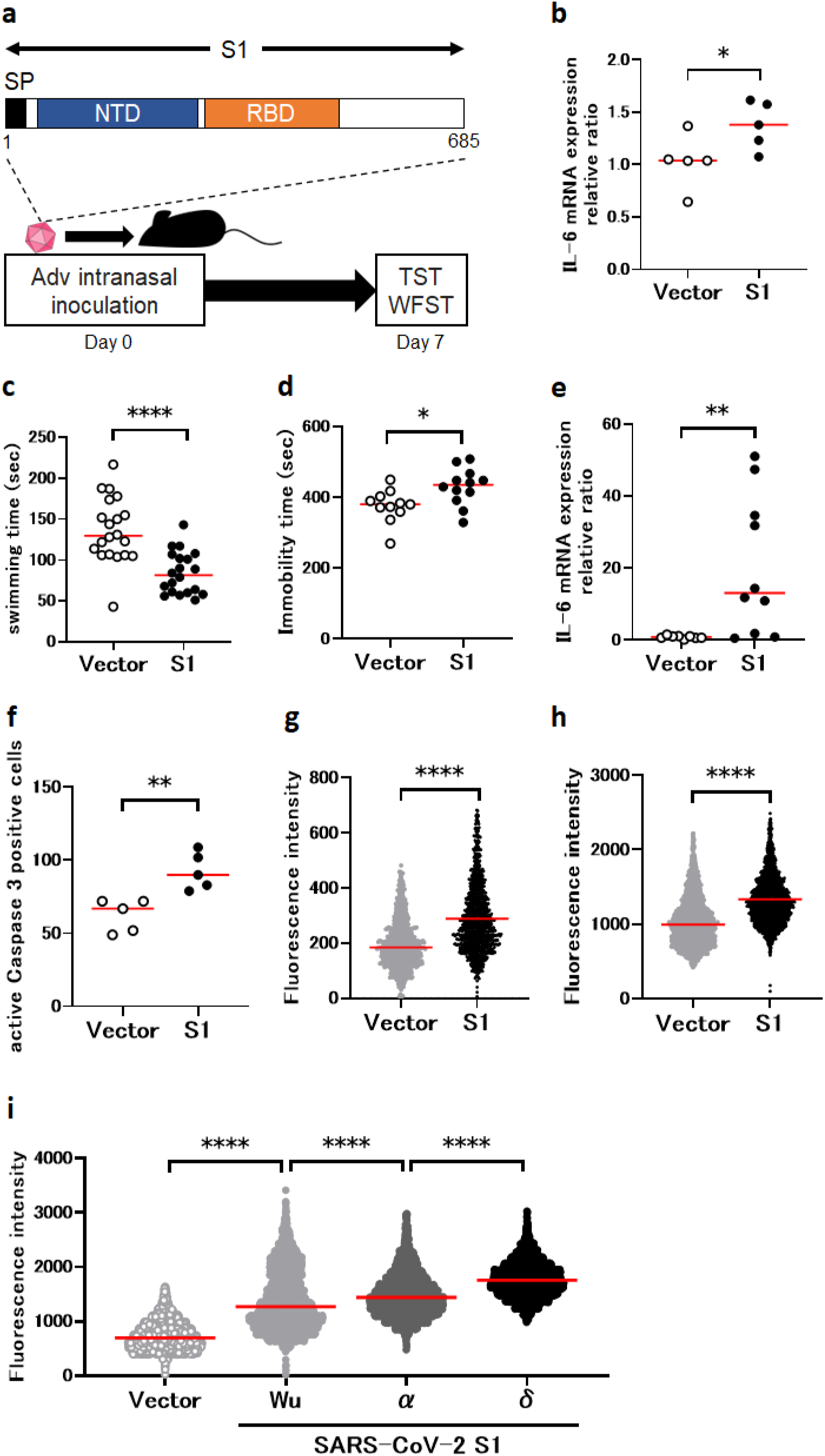
Creation of COVID-19 encephalopathy model using S1 protein and analysis of S1 protein function. **a**, S1 mouse creation and evaluation **b**, Enhanced IL-6 expression in S1 mouse brain (vector control, n=5; S1 mouse, n=5; Mann-Whitney U-test; median values; *, p < 0.05). **c**, Reduced swimming time in S1 mouse WFST; i.e., increase in fatigue (vector control, n=20; S1 mouse, n=20; Mann-Whitney U-test; median values; ****, p < 0.0001). **d**, Prolonged immobility time in S1 mouse TST, i.e., increase in depressive symptoms (vector control, n=11; S1 mouse, n=12; Mann-Whitney U-test; median values; *, p < 0.05). **e**, Enhanced IL-6 expression in S1 mouse lung (vector control, n=10; S1 mouse, n=10; Mann-Whitney U-test; median values; **, p < 0.01). **f**, Measurement of apoptosis induction in S1 mouse olfactory bulb based on caspase 3 positive cell count (vector control, n=5; S1 mouse, n=5; Mann-Whitney U-test; median values; **, p < 0.01). **g**, Measurement of intracellular calcium concentration in human A549 cells using Fluo 4 fluorescence intensity (vector control, n=828; S1 Adv, n=1085; Mann-Whitney U-test; median values; ****, p < 0.0001). **h**, Measurement of intracellular calcium concentration in mouse 3T3 cells (vector control, n=2268; S1 Adv, n=1830; Mann-Whitney U-test; median values; ****, p < 0.0001). **i**, Comparison of intracellular calcium concentration elevation among Wu strain, α strain and δ strain (vector control, n=3562; Wu strain, n=2711; α strain, n=2837; δ strain, n=2339; Kruskal-Wallis test followed by Dunn’s post hoc test; median values; ****, p < 0.0001)

In the lungs, a correlation of inflammatory cytokine production with S1 mRNA expression (Extended Data Fig.1c, d), suggested that inflammation in the lungs was due to the inflammation-inducing action of S1 protein ^23^. In contrast, since S1 mRNA expression was not detected in the brain, this suggested that inflammation in the brain was due to an indirect effect (Extended Data Fig.1e).

### Analysis of S1 protein function

When examining pathological changes in the brain to elucidate the mechanism for the inflammation promoting mechanism of S1 protein, we observed apoptosis induction in the olfactory bulb of the S1 mouse (Fig. 1f). In previous research, we demonstrated that human herpesvirus-6 (HHV-6) protein produced in the olfactory epithelium and olfactory bulb sustentacular cells induced apoptosis in the olfactory bulb causing depression, and this was due to its calcium elevating activity^24^. Observing such activity for S1 protein as well (Fig. 1g, h), we considered that the apoptosis in the S1 mouse olfactory bulb and depressive symptoms were due to the calcium elevating activity of S1 protein.

To examine the involvement of S1 protein’s intracellular calcium increasing activity in pathogenicity, we measured this for S1 protein from the Wu strain, α strain and δ strain. The strength of intracellular calcium concentration elevation activity in cells expressing the different strains was in the order: Wu<α<δ (Fig. 1i). This was consistent with the order for the severity and mortality rates of these strains: Wu<α<δ ^25,26^.

### Elucidation of inflammation enhancing mechanism

To elucidate a mechanism for the effect of olfactory bulb damage on the brain and the body overall, we focused on neurotransmitters in neurons connecting to the olfactory bulb. There was a decrease in cells positive for the acetylcholine synthesizing enzyme choline acetyltransferase (ChAT) in the medial septal (MS) and diagonal band of Broca (DBB) (Fig. 2a, b) and the acetylcholine level in the brain overall was also decreased (Fig. 2c). We therefore examined a relationship between brain inflammation in the S1 mouse and the anti-inflammatory response in which acetylcholine plays a role, called cholinergic anti-inflammatory pathway (CAP) ^9^. Since an α7 nicotinic acetylcholine receptor (α7nAchR) agonist was found to activate CAP ^27^, one week after inoculation with S1 Adv, we administered the α7nAchR agonist PNU282987 intracerebroventricularly (i.c.v.) and after an hour, measured inflammatory cytokine expression levels in the brain (Fig. 2d). This resulted in normalizing the enhanced inflammatory cytokine expression in the S1 mouse (Fig. 2e). These observations suggest that brain inflammation in the S1 mouse resulted from CAP disruption.

**Fig. 2:**
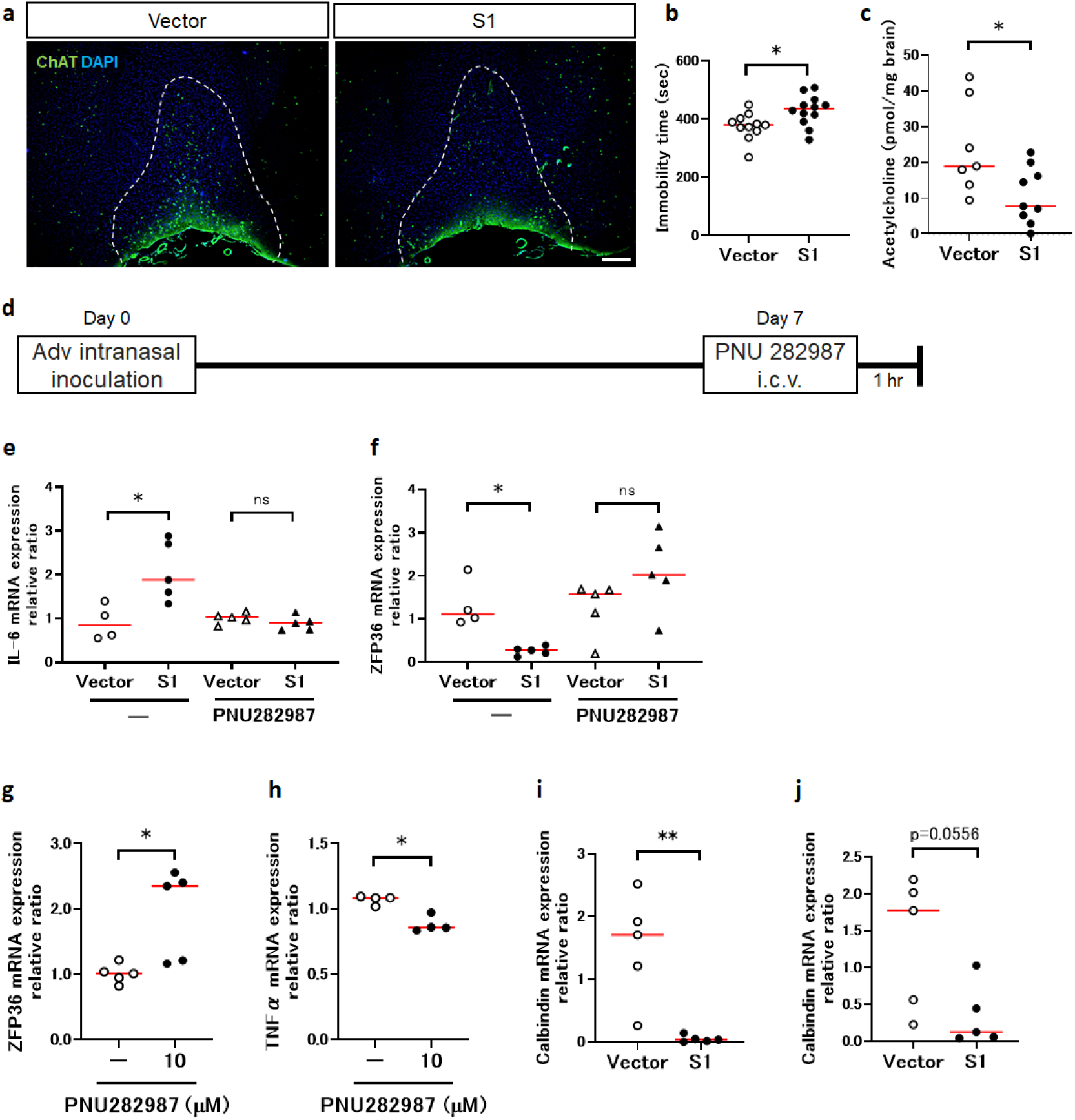
Disruption of cholinergic anti-inflammatory pathway (CAP) in S1 mouse brain. **a**, Decreased ChAT positive cells in S1 mouse MS and DBB (green, ChAT; blue, DAPI; scale bar, 200 μm). **b**, Decreased ChAT positive cells in S1 mouse MS and DBB (vector control, n=5 S1 mouse, n=5; Mann-Whitney U-test; median values; *, p < 0.05). **c**, Intracerebral acetylcholine (vector control, n=8; S1 mouse, n=9; Mann-Whitney U-test; median values; *, p < 0.05). **d**, Adv inoculation, PNU282987 administration, brain harvest schedule. **e**, Enhanced IL-6 expression in S1 mouse brain and effect of PNU282987 (no treatment; vector control, n=4 S1 mouse, n=5; Mann-Whitney U-test; median values; *, p < 0.05. PNU282987; vector control, n=5 S1 mouse, n=5; Mann-Whitney U-test; median values; ns, not significant.). **f**, Decreased ZFP36 expression in S1 mouse brain and effect of PNU282987 (no treatment; vector control, n=4 S1 mouse, n=5; Mann-Whitney U-test; median values; *, p < 0.05. PNU282987; vector control, n=5 S1 mouse, n=5; Mann-Whitney U-test; median values; ns, not significant.). **g**, ZFP36 mRNA expression in U373 cells (LPS stimulation, n=5; LPS stimulation+PNU282987 treatment, n=5; Mann-Whitney U-test; median values; *, p < 0.05). **h**, TNF *α* mRNA expression in U373 cells (LPS Stimulation, n=5; LPS stimulation+PNU282987 treatment, n=5; Mann-Whitney U-test; median values; *, p < 0.05). **i**, Expression of calbindin in olfactory bulb of S1 mice additionally administered LPS (vector control, n=5 S1 mouse, n=5; Mann-Whitney U-test; median values; **, p < 0.01). **j**, Expression of calbindin in brain of S1 mouse additionally administered LPS (vector control, n=4 S1 mouse, n=5; Mann-Whitney U-test; median values.)

It has been known for some time that nerves in the CAP suppress inflammation ^9,28^ but inflammation suppressing factors were unknown. In the S1 mouse, the expression of the mRNA binding protein ZFP36 ^10^, which has an inflammatory cytokine mRNA degrading function, was reduced when inflammation was present in the brain and was increased by PNU282987 (Fig. 2f). Also, administering PNU282987 to the human astrocyte cell line U373 ^29^ expressing α7nAchR enhanced ZFP36 expression and suppressed TNF *α* expression (Fig. 2g, h). These findings suggest that ZFP36 functions as an inflammation-suppressing factor in the CAP.

In order to investigate the influence of systemic inflammation in brain inflammation, we administered lipopolysaccharide (LPS) intraperitoneally to the S1 mouse to enhance systemic inflammation and investigated expression of neural differentiation markers in the olfactory bulb and brain. As a result, the expression of calbindin ^30,31^, a GABAergic neuron marker, was markedly decreased. (Fig. 2i, j). As no changes were observed in other neuron differentiation markers (Extended Data Fig. 2a, b), this observation was considered specific to GABAergic neurons.

### Improvement in brain symptoms with administration of donepezil

The acetylcholine esterase inhibitor donepezil is a central cholinergic agent used clinically for the treatment of dementia ^32^. To investigate the possibility of its repurposing for the treatment of neurological complications in COVID-19, we investigated symptom improvement due to donepezil in the S1 mouse.

We administered donepezil at the usual dose for animal experiments (4 mg/kg/day) from the day of S1 Adv inoculation daily in drinking water (Fig. 3a). Administration of donepezil resulted in normalizing inflammatory cytokines (IL-6, TNFα) that had been enhanced by S1 protein (Fig. 3b, Extended Data Fig. 3a). Since IL-6 and TNFα genes are targets of ZFP36 ^10,33^, this finding suggests that donepezil brought about a recovery in CAP. However, administration of donepezil did not mitigate the enhanced inflammatory cytokine production in the lungs (Extended Data Fig. 3b, c, d).

**Fig. 3:**
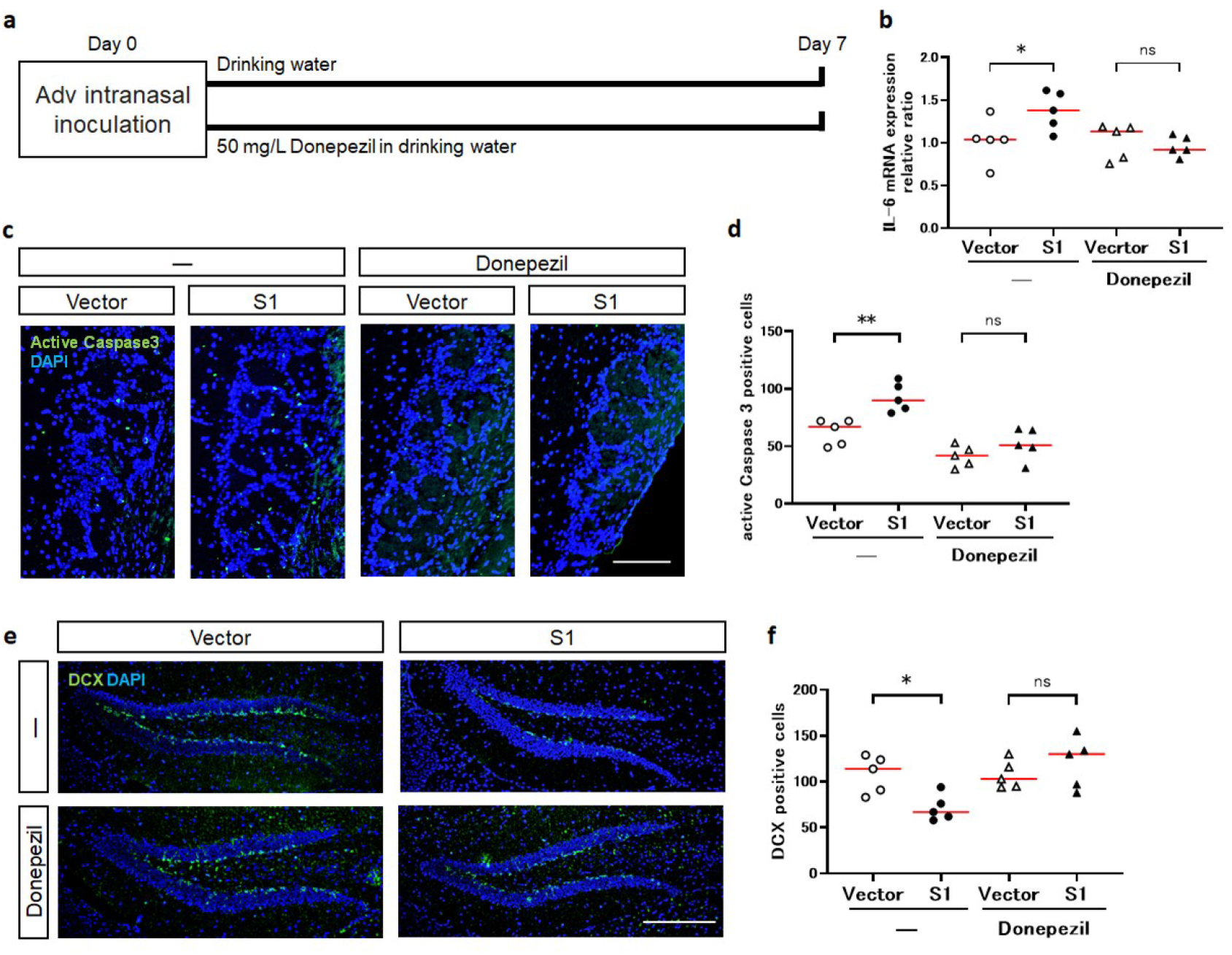
Mitigation of olfactory and brain dysfunction in S1 mouse due to donepezil. **a**, Adv inoculation, donepezil administration, brain harvest schedules. **b**, Enhanced IL-6 expression in S1 mouse brain and improvement due to donepezil (no treatment; vector control, n=4, S1 mouse, n=5; Mann-Whitney U-test; median values; *, p < 0.05. Donepezil; vector control, n=5, S1 mouse, n=5; Mann-Whitney U-test; median values; ns, not significant). **c**, Decreased apoptosis cells in olfactory bulb of S1 mouse and improvement due to donepezil (green, Active caspase 3; blue, DAPI; scale bar, 100μm). **d**, Increased caspase 3 positive apoptotic cells in S1 mouse olfactory bulb and improvement due to donepezil (no treatment; vector control, n=5, S1 mouse, n=5; Mann-Whitney U-test; median values; **, p < 0.01. Donepezil; vector control, n=5 S1 mouse, n=5; Mann-Whitney U-test; median values; ns, not significant). **e**, Decreased DCX positive cells in S1 mouse hippocampus and improvement due to donepezil (green, DCX; blue, DAPI; scale bar, 200μm). **f**, Decreased DCX positive cells in S1 mouse hippocampus and improvement due to donepezil (no treatment; vector control, n=5, S1 mouse, n=5; Mann-Whitney U-test; median values; *, p < 0.05. Donepezil; vector control, n=5 S1 mouse, n=5; Mann-Whitney U-test; median values; ns, not significant.).

Regarding pathological changes in the brain, donepezil mitigated the enhanced apoptosis in the olfactory bulb of the S1 mouse (Fig. 3c, d). There was also improvement due to donepezil with regard to doublecortin (DCX) positive cells, which indicate hippocampal neurogenesis and had been decreased in the S1 mouse (Fig. 3e, f). A decrease in hippocampal neurogenesis has been reported to be associated with depressive symptoms ^34^ and memory impairment ^35^, so we considered it likely that donepezil would improve depressive symptoms and memory impairment in COVID-19. However, donepezil did not reverse a decrease in ChAT positive cells in the MS and DBB (Extended Data Fig. 3e, f).

### Treatment of COVID-19 with donepezil

With the objective of using donepezil clinically to treat neurological complications in COVID-19 in mind, we examined the drug’s effects on symptoms as well as the dose that would be effective.

We administered donepezil at the usual dose for animal experiments (4 mg/kg/day) daily in drinking water from the day of S1 Adv inoculation (Fig. 4a). The administration of donepezil mitigated the increased fatigue determined in the weigh-loaded swimming test and depressive symptoms determined from the tail suspension test (Fig. 4b, c).

**Fig. 4:**
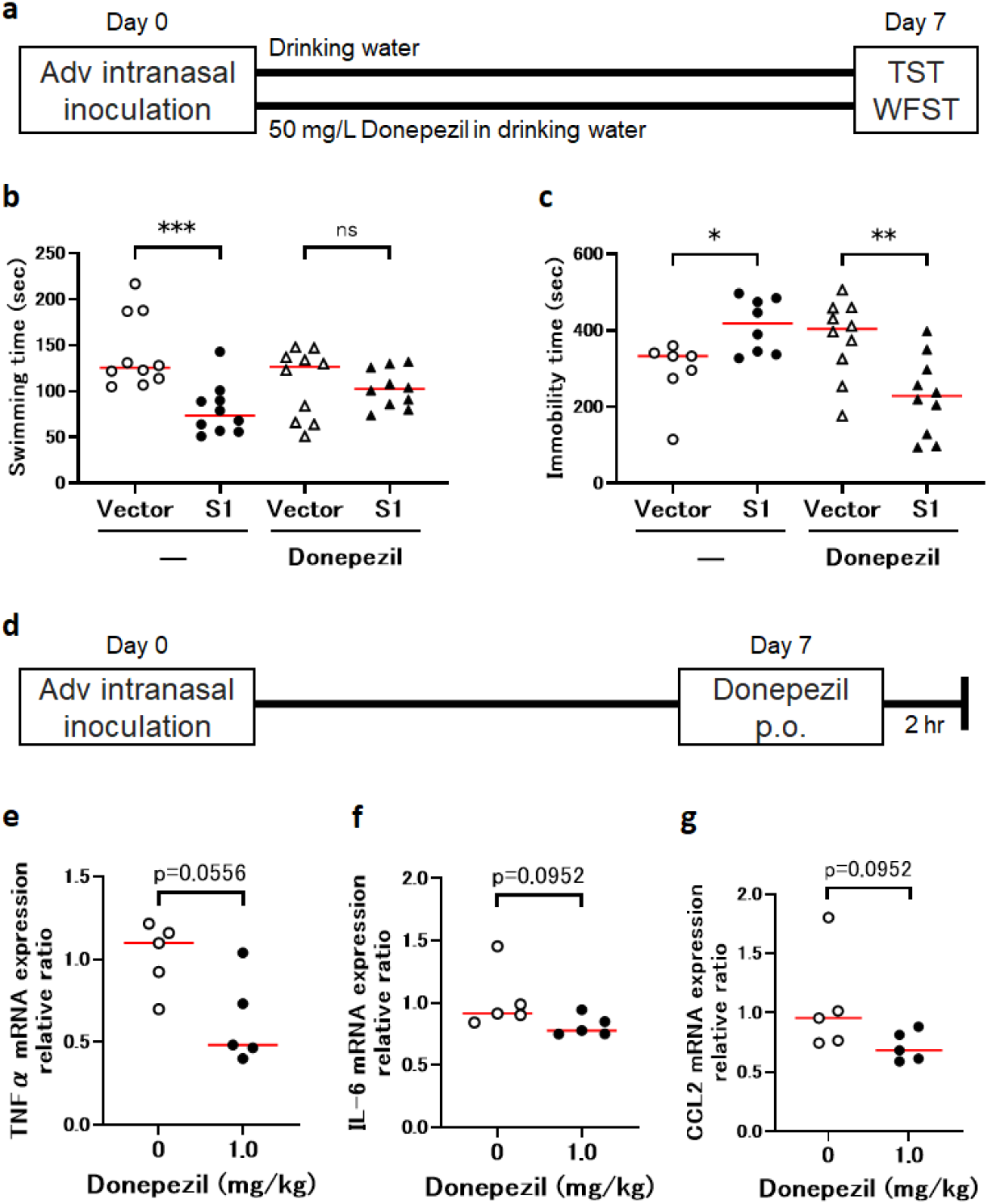
Therapeutic effects of donepezil with respect to COVID-19. **a**, Adv inoculation, donepezil administration, behavior experiment schedules. **b**, Decreased S1 mouse swimming time in WFST, i.e., increase in fatigue, and therapeutic effect of donepezil (no treatment; vector control, n=10, S1 mouse, n=10; Mann-Whitney U-test; median values; ***, p < 0.001. Donepezil; vector control, n=10 S1 mouse, n=10; Mann-Whitney U-test; median values; ns, not significant). **c**, Increased immobility time in S1 mouse TST and therapeutic effect of donepezil (no treatment; vector control, n=7, S1 mouse, n=8; Mann-Whitney U-test; median values; *, p < 0.05. Donepezil; vector control, n=10 S1 mouse, n=10; Mann-Whitney U-test; median values; ns, not significant). **d**, Adv inoculation, donepezil administration, organ harvest schedules **e**, Inhibitory effect of donepezil on TNFα expression in S1 mouse brain (0mg/kg, n=5; 1mg/kg, n=5; Mann-Whitney U-test; median values). **f**, Inhibitory effect of donepezil on IL-6 expression in S1 mouse brain (0mg/kg, n=5; 1mg/kg, n=5; Mann-Whitney U-test; median values). **g**, Inhibitory effect of donepezil on CCL2 expression in S1 mouse brain (0mg/kg, n=5; 1mg/kg, n=5; Mann-Whitney U-test; median values).

Next, we investigated whether the normal dose of donepezil used for dementia would be effective. We converted the normally administered dose for dementia patients (5 mg/day) for mice (1 mg/kg/day) and administered a single dose orally. Administration was one week after inoculation of S1 Adv and measurements were made 2 hr. later (Fig. 4d). A tendency toward mitigation of the enhanced expression of inflammatory cytokines (IL-6, TNFα) and a chemokine (CCL-2) was observed in the S1 mouse brain (Fig. 4e, f, g). This result indicates the possibility of donepezil being effective against brain inflammation in COVID-19 at the normal dose.

## Discussion

Through the inoculation of a non-proliferative adenovirus vector expressing S1 protein into the nasal cavity, we created an S1 mouse exhibiting neurological complications such as brain inflammation, fatigue and depressive symptoms, and inflammation in the lungs.

Since inflammatory cytokine and S1 protein mRNA quantities in the lungs of the S1 mouse were correlated, we considered the inflammation to be due to an inflammation-inducing action of S1 protein ^21,36 23^. This suggests that pneumonia due to infection by SARS-CoV-2 similar to that in COVID-19 had been reproduced in the S1 mouse. In addition, a lack of S1 mRNA expression in the brain led us to conclude that we had reproduced a state of encephalopathy similar to that reported for COVID-19, where the inflammation is not due to viral proliferation in the brain but caused indirectly.

In the brain of the S1 mouse, we observed a decrease in cells positive for ChAT in the MS and DBB as well as a decrease in the amount of acetylcholine in the brain and CAP disruption ^9,28^. We considered neurological complications had developed in the S1 mouse because inflammation in the brain caused by inflammatory cytokines produced in the lungs could not be suppressed due to CAP disruption. Since it was reported that COVID-19 symptoms were mitigated by nicotine administered through smoking, much attention has been given to a relationship between COVID-19 and CAP ^37,38^. Decreased expression of acetylcholine receptors in the peripheral blood of COVID-19 patients has also been reported ^39^.

Systemic inflammation due to proliferation of the virus throughout the body in COVID-19 could be stronger than that in observed in the S1 mouse. Therefore, we enhanced systemic inflammation through intraperitoneal administration of lipopolysaccharide (LPS), which resulted in markedly reduced expression of the GABAergic marker calbindin ^32^ in the olfactory bulb and the brain. The olfactory bulb has many GABAergic neurons ^40^ and it has been reported that GABAergic dysfunction is the cause of brain fog in long COVID ^41,42^. Thus, our finding may help explain the pathogenesis mechanism of olfactory damage and brain fog in long COVID.

Since it was reported that surgically-induced disruption of olfactory bulb function through olfactory bulbectomy (OBX) resulted in a decrease in ChAT positive cells in the MS^43^, the decrease in ChAT positive cells and disruption of the CAP in the S1 mouse could be due to olfactory bulb dysfunction. Thus, severe damage to the olfactory bulb due to SARS-CoV-2 infecting the olfactory system^44,45^ could be the main cause of CAP disruption in COVID-19. However, the olfactory bulb damage in the S1 mouse was thought to be due to the calcium elevating activity of S1 protein. Although the influence of calcium elevating activity in olfactory bulb damage in COVID-19 is unknown, our finding that strength of calcium elevating activity for different strains was in the order Wu<α<δ is of great interest because this is consistent with the order of their severity and mortality rates ^25,26^. Regarding the calcium-related pathogenicity of spike protein and SARS-CoV-2, an association with syncytia formation has been found ^46^. As it has been reported that the α strain and δ strain have the same ability to form syncytia ^47^, the mechanism indicated in the present study may make consistency between strength of calcium enhancing activity and pathogenicity easier to explain.

OBX is also known to cause depressive-like symptoms ^48^ and intracerebral inflammation ^49^, in addition to CAP disruption. This suggests that the brain inflammation and neurological complications seen in COVID-19 patients are due to the disruption of the CAP caused by acetylcholine deficiency resulting from olfactory bulb destruction.

To mitigate the pathological state in the S1 mouse, we administered donepezil for inhibition of choline esterase, the enzyme that breaks down acetylcholine. This resulted in reversing the inflammation, reduced hippocampal neuro-genesis and apoptosis in the olfactory bulb, and the fatigue and depressive symptoms associated with them also disappeared. As donepezil readily passes through the blood-brain barrier (BBB) and has a brain to blood drug concentration ratio of 4 or above ^50^, we considered that these improvements were due to the action of donepezil acting in the brain.

On the other hand, the decrease in ChAT positive cells in the MS and DBB was not reversed by donepezil. This finding again suggests that a decrease in ChAT positive cells is positioned upstream of a series of pathological states and is considered to provide a basis for an increase in acetylcholine due to donepezil having a therapeutic effect on the overall symptoms in the S1 mouse.

Our findings indicate that administering donepezil according to the same administration and dosage directions as for the treatment of dementia would suppress brain inflammation. This suggests that within the guaranteed safety range, donepezil could be applied for the treatment of brain inflammation and neurological complications in COVID-19 and that drug repositioning would allow it to be used in clinical practice at an early date. As many aspects of their pathogenesis mechanisms are unknown and effective treatments have yet to be established, the results of the present study should not only contribute to the elucidation of pathologies, but also have high clinical value.

## Limitation

The effects of donepezil in the present research were obtained in an animal study. Therefore, in order to use it in the treatment of COVID-19 sequelae, its effects on patients will need to be confirmed through clinical trials or other methods.

## Acknowledgments

We thank Mr. Alexander Cox for editorial assistance with the manuscript. This research was supported by AMED under Grant Number JP20fk0108541.

## Author Contributions

N.O. and K.K. contributed to study design. N.O., K.S., A.I., N.K., and K.K. contributed to data collection or interpretation. K.K. coordinated all experiments.

## Methods

### Animals

Male 8-week-old C57BL/6NCrSlc mice obtained from Sankyo laboratories were used for all experiments. All mice were housed under standard conditions (12-hour light–dark cycle [lights-on at 8:00 a.m.] at 24 ± 1 °C) with food and water provided ad libitum. All animal experiments were performed in accordance with animal experiment regulations and approved by the Animal Care and Use Committee of the Jikei University School of Medicine.

### Viruses and cells

The mouse fibroblast-like cell line 3T3 and human lung epithelial cell line A549 were cultured in Dulbecco’s modified Eagle’s medium (DMEM) containing 10% FBS. The recombinant adenovirus was produced using an Adenovirus Dual Expression Kit (Takara Bio) in accordance with the manufacturer’s protocol. The SARS CoV-2 genes were cloned into an adenovirus cosmid vector (pAxCAwtit2) using standard methods (SARS CoV-2 genes/pAxCAwtit2). HEK293 cells were transfected with the SARS CoV-2 genes/pAxCAwtit2 cosmid and a cosmid that did not contain the target gene (pAxcwit2). The recombinant adenovirus was prepared in HEK293 cells and purified with an Adeno-X Virus Purification kit (Takara Bio). The purified virus titer was determined using an Adeno-X Rapid Titer kit (Takara Bio). Sex and age at sampling for the cells were as follows: 3T3, male embryo; A549, male 58Y; HEK293, female fetus.

### Measurement of intracellular calcium concentration

3T3 cells or A549 cells were cultured on a 96-well plate and transient overexpression of each protein -S1 (Wu, *α* and *δ*), NTD and Spike - was achieved for 24 h using ProFection Mammalian Transfection System (Promega) or adenovirus infection. Subsequently, cells were induced to take up Fluo 4-AM using Calcium Kit II-Fluo 4 (Dojindo) and then the fluorescence intensity in individual cells was measured using an ArrayScan XTI instrument (Thermo Fisher).

### Nasal inoculation of adenovirus vectors

For the nasal inoculation of adenovirus vectors, 8-week-old male C57BL/6 mice were anaesthetized with isoflurane. Recombinant adenoviruses were diluted in sterile water (not in isotonic buffer). A drop (25 μL) of S1-Ad or Vector-Ad solution containing the virus at a titer of 1 × 10^9^ infectious units (ifu) /mL was placed at the entrance of the nasal cavity of the mouse. The solution entered the cavity through spontaneous respiration.

### Drugs

The cholinesterase inhibitor donepezil (FUJIFILM Wako) was administered via the drinking water (40 mg/L, 5mg/kg/day) from the day of intranasal inoculation. For oral administration, donepezil was diluted in saline and administered to mice by feeding tube at 0, 1 and 2 mg/kg. The α7nAChR agonist PNU282987 (FUJIFILM Wako) was diluted in saline and administered intracerebroventricularly at 400 nmol/mouse. Lipopolysaccharide (LPS) from Escherichia coli O111:B4 (Merck) was diluted in saline and administered intraperitoneally to S1 mice at 5 mg/kg. The brain and olfactory bulb were removed 15 and 30 minutes after administration, respectively, and used for gene expression analysis.For all oral, intracerebroventricular and intraperitoneal administrations, control mice received the solvent saline.

### Animal behaviour tests

Seven days after the inoculation of S1-Ad or Vector-Ad, a tail suspension test (TST) and a weight-loaded forced swim test (WFST)were performed to assess depression- and fatigue-like behaviour. In the TST, the period of immobility during 10 min was analysed by TailSuspScan software (CleverSys Inc). In the WFST, a weight equivalent to 10% of body weight was attached to the root of the mouse tail, the time taken for the mouse’s nose to be below the surface of the water for 10 seconds was measured by TopScan software (CleverSys Inc). At 24 h, after these behaviour tests, the mice were euthanized for real-time PCR and immunohistochemical staining.

### Real-time PCR

Total RNA was purified from animal tissues using the RNeasy Mini QIAcube Kit (Qiagen). cDNA was synthesized from total RNA with a PrimeScript RT Reagent Kit (Takara Bio). mRNA amounts were quantified with Premix Ex Taq (Perfect Real Time) (Takara Bio), and the Applied Biosystems 7300 Real-Time PCR system (Thermo Fisher). The thermal profile was 95 °C for 30 s, followed by 45 cycles of 95 °C for 5 s and 60 °C for 31 s. Data analysis was performed with Sequence Detection Software version 1.4 (Thermo Fisher). Mouse IL-6, mouse IL-1β, mouse TNFα, mouse CCL-2, mouse ZFP36, mouse Calbindin, mouse GFAP, mouse Nestin, mouse IL-17a, human 18S and SARS-CoV-2 S1 were measured using the primers and probes described in Supplementary Table 1.

### Immunohistochemistry

For the staining of tissues, mice were sacrificed and transcardially perfused with saline, followed by 10% neutral buffered formalin (pH 7.4). Formalin-fixed tissues were then embedded in paraffin and sectioned. For immunofluorescence staining, paraffin-embedded sections were deparaffinized, and blocking was performed in Image-iT FX Signal Enhancer (Life Technologies) for 30 min. Primary antibodies were obtained from Abcam (active caspase-3, doublecortin and Choline Acetyltransferase). Secondary antibodies were obtained from Thermo (Alexa Fluor 488 Goat Anti-rabbit IgG (H+L)). The antibodies were diluted with Can Get Signal Immunostain Solution A (TOYOBO). After the samples had been mounted on a slide with a cover glass, they were observed under a Keyence BZ-9000 fluorescence microscope.

### Measurement of acetylcholine levels in mouse brain

Seven days after the inoculation of S1-Ad or Vector-Ad, mice were sacrificed and their brains immediately harvested. The amount of acetylcholine in the brain was measured by the Choline/Acetylcholine Assay Kit (abcam). Fluorescence was measured by TriStar LB941 (Berthold).

### Expression of anti-inflammatory factors through stimulation of *α* 7nicotinic acetylcholine receptors

U373 cells were cultured on a 6-well plate and stimulated with 10 µM PNU282987. After culturing U373 cells in the presence of 10µg/mL LPS (Merck) for 48 hr, they were stimulated with 10 µM PNU282987.Cells were recovered 2hr after stimulation followed by RNA purification and cDNA synthesis by the methods mentioned above and then expression of anti-inflammatory factors was analyzed by RT-qPCR. Human IL-6, Human ZFP36 and human 18S were measured using the primers and probes described in Supplementary Table 1.

### Quantification and statistical analysis

The Shapiro-Wilk normality test was performed to assess the normality of distributions. To compare two different groups, the Mann–Whitney U-test was used as the nonparametric test. To compare multiple groups, the Kruskal-Wallis test, and then Dunn’s post hoc test, were used as non-parametric tests.Red horizontal lines are medians. *P* < 0.05 was considered significant. Spearman’s rank correlation coefficients were used to determine correlations between variables. Statistical analyses were performed with Prism 8 (GraphPad) and BellCurve for Excel (Social Survey Research Information Co., Ltd.).

**Table 1:**
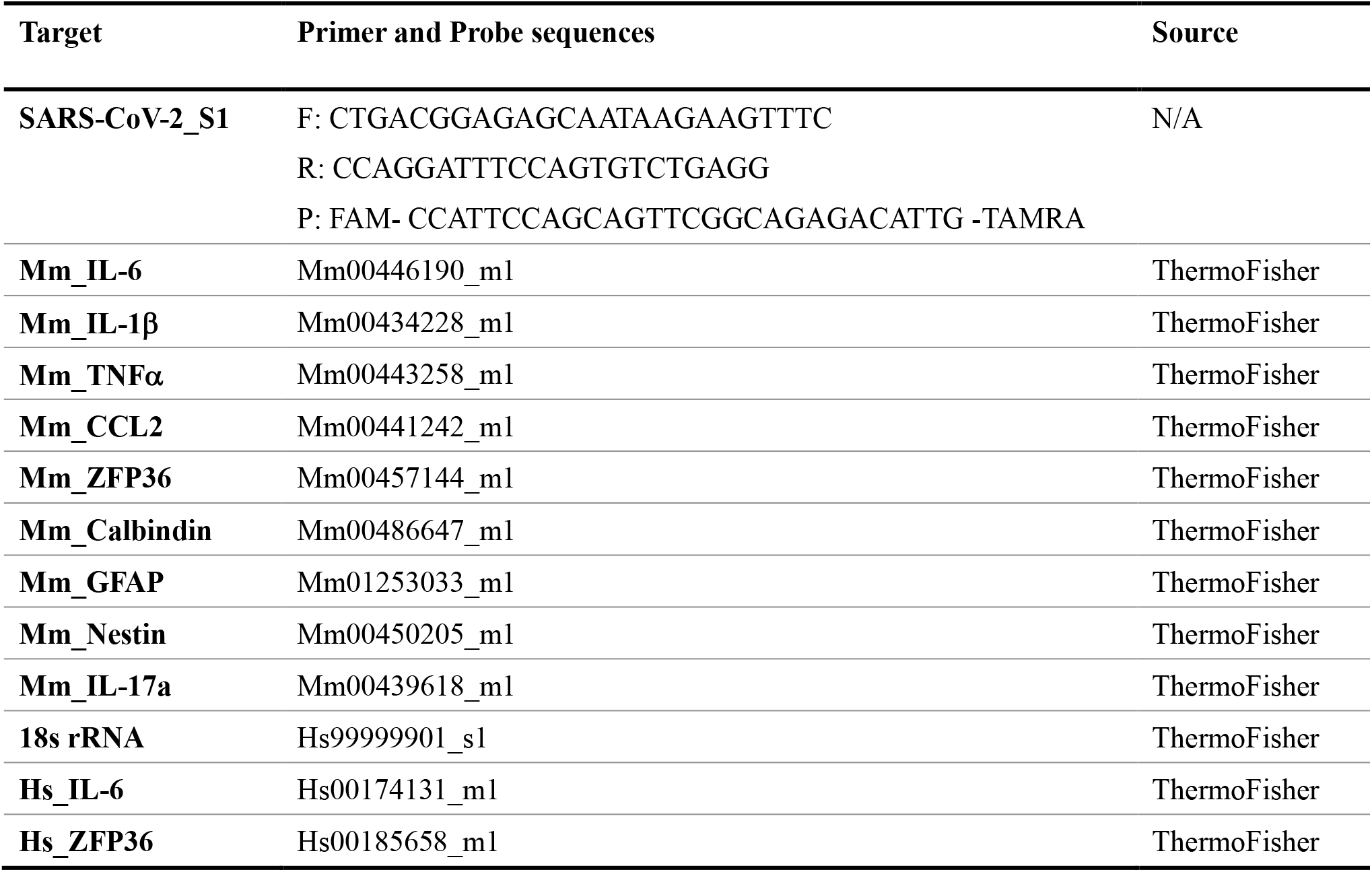
List of primer and probe sequences used in this study.

