## Supplemental data for "SARS-CoV-2 causes brain inflammation via impaired neuro-immune interactions"

Extended Data Fig. 1: Creation of COVID-19 encephalopathy model using S1 protein and analysis of S1 protein function

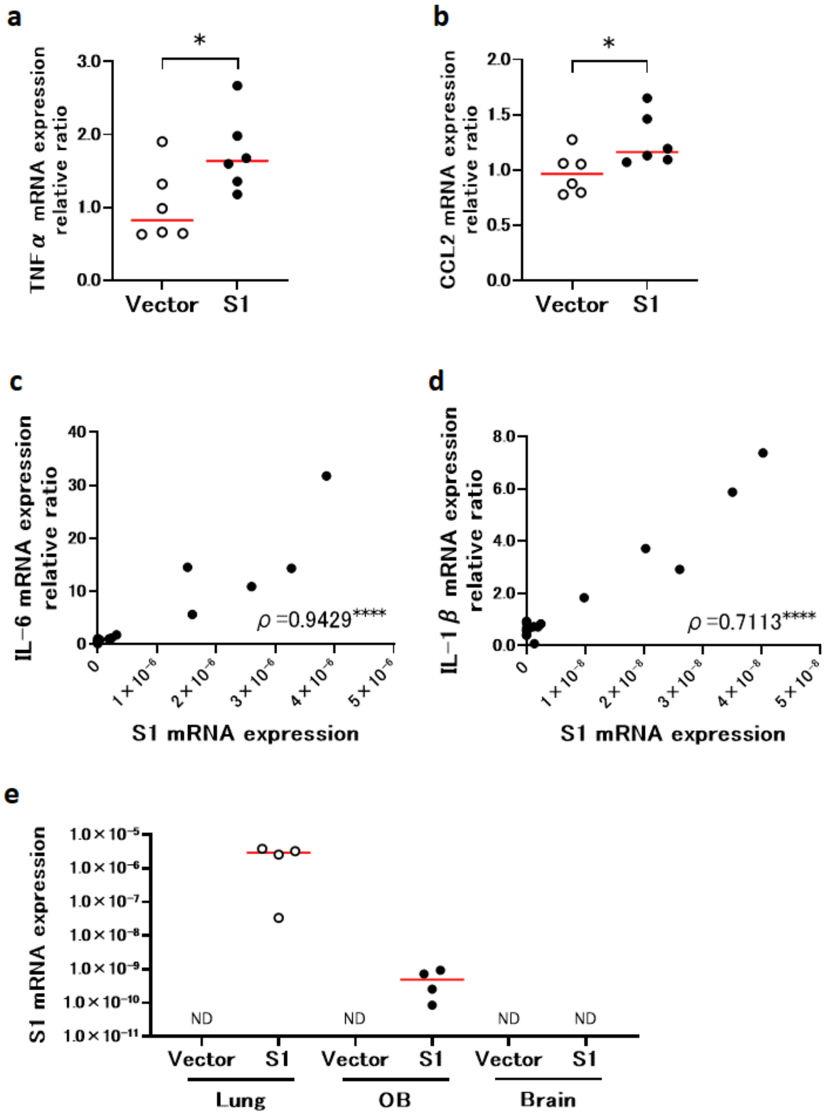


**Extended Data Fig. 1: Creation of COVID-19 encephalopathy model using S1 protein and analysis of S1 protein function**

**a**, Enhanced TNFα expression in S1 mouse brain (vector control, n=6; S1 mouse, n=6; Mann-Whitney U-test; median values; ∗, $p < 0.05$). **b**, Enhanced CCL-2 expression in S1 mouse brain (vector control, n=6; S1 mouse, n=6; Mann-Whitney U-test; median values; ∗, $p < 0.05$). **c**, Correlation of S1 mRNA expression and IL-6 mRNA expression in lung (n=22; Spearman's rank correlation test; r =0.9429, ∗∗∗∗, $p < 0.0001$). **d**, Correlation of S1 ｍRNA expression and IL-1β mRNA expression in lung (n=18; Spearman's rank correlation test; r =0.7113, ∗∗∗∗, $p < 0.0001$) . **e**, S1 mRNA expression in lung, olfactory bulb and brain（Ratios of S1 to 18s ribosomal RNA are shown.）(vector control, n=4 each; S1 mouse, n=4 each median values).

Extended Data Fig. 2: Changes in nerve differentiation marker expression in S1 mice additionally administered LPS

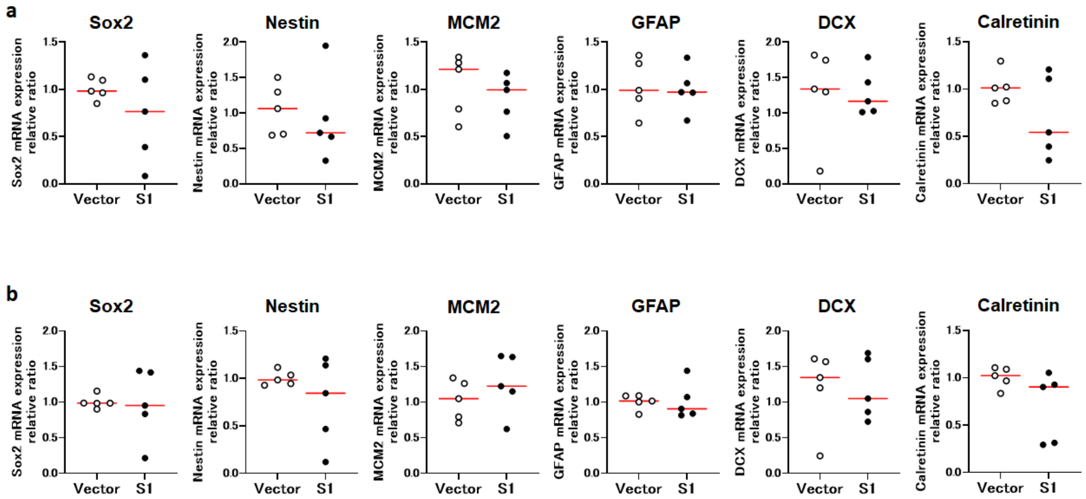


**Extended Data Fig. 2: Changes in nerve differentiation marker expression in S1 mice additionally administered LPS**

**a,** Changes in expression of indicated nerve differentiation marker genes in olfactory bulb of S1 mice additionally administered LPS. **b**, Changes in expression of indicated nerve differentiation marker genes in brains of S1 mice additionally administered LPS.

Extended Data Fig. 3: Mitigation of olfactory and brain dysfunction in S1 mouse due to donepezil

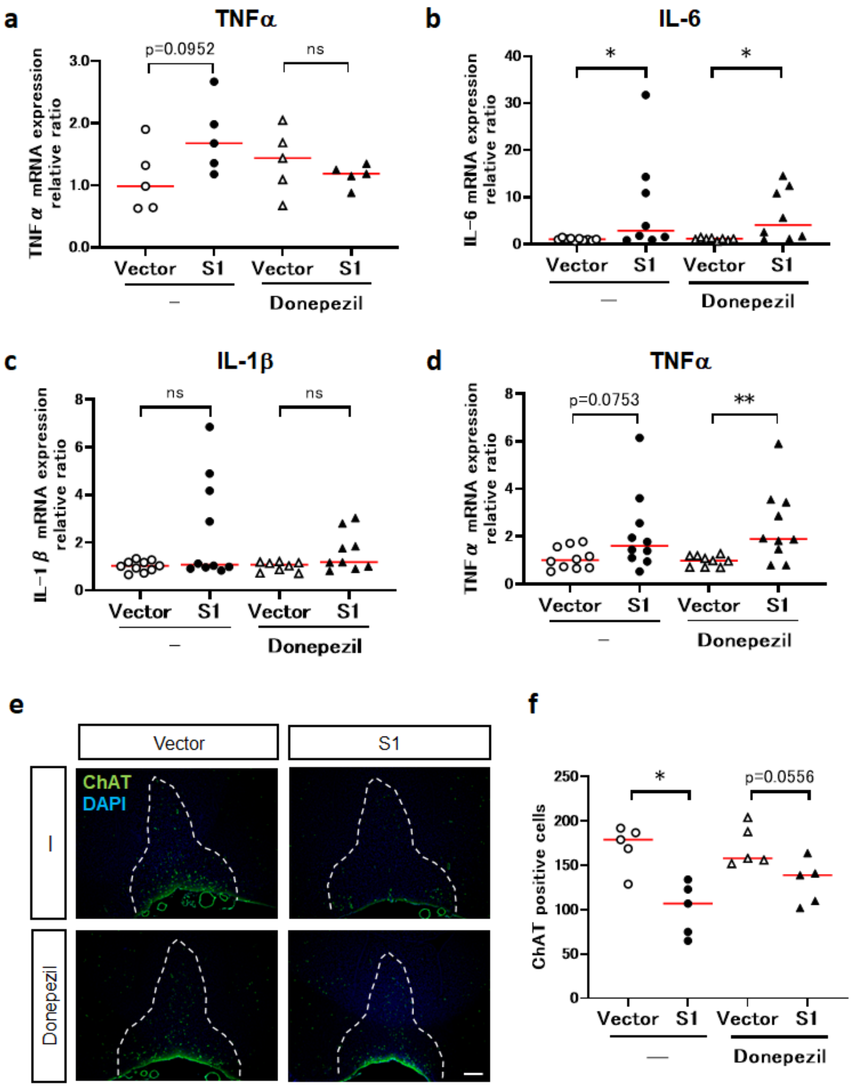


**Extended Data Fig. 3: Mitigation of olfactory and brain dysfunction in S1 mouse due to donepezil**

**a**, Enhanced TNFαexpression in S1 mouse brain and improvement due to donepezil (no treatment; vector control, n=5, S1 mouse, n=5; Mann-Whitney U-test; median values. Donepezil; vector control, n=5, S1 mouse, n=5; Mann-Whitney U-test; median values; ns, not significant). **b**, Enhanced IL-6 expression in S1 mouse lung and effect of administering donepezil (no treatment; vector control, n=8, S1 mouse, n=8; Mann-Whitney U-test; median values; ∗, $p < 0.05$. Donepezil; vector control, n=8, S1 mouse, n=8; Mann-Whitney U-test; median values; ∗, $p < 0.05$). **c,** Enhanced IL-1βexpression in S1 mouse lung and effect of administering donepezil (no treatment; vector control, n=10, S1 mouse, n=10; Mann-Whitney U-test; median values; ns, not significant. Donepezil; vector control, n=8, S1 mouse, n=8; Mann-Whitney U-test; median values; ns, not significant). **d,** Enhanced TNFαexpression in S1 mouse lung and effect of administering donepezil (no treatment; vector control, n=10, S1 mouse, n=10; Mann-Whitney U-test; median values. Donepezil; vector control, n=10, S1 mouse, n=10; Mann-Whitney U-test; median values; ∗∗, $p < 0.01$). **e**, Decreased ChaT positive cells in S1 mouse MS and DBB and effect of donepezil (green, DCX; blue, DAPI; scale bar, 200μm). **f**, Decreased ChAT positive cells in S1 mouse MS and DBB and effect of donepezil (no treatment; vector control, n=5, S1 mouse, n=5; Mann-Whitney U-test; median values; ∗, $p < 0.05$. Donepezil; vector control, n=5 S1 mouse, n=5; Mann-Whitney U-test; median values).
